# Microtubule inner proteins of *Plasmodium* are essential for transmission of malaria parasites

**DOI:** 10.1101/2023.10.19.562943

**Authors:** Franziska Hentzschel, Annika M. Binder, Lilian P Dorner, Lea Herzel, Fenja Nuglish, Meslo Sema, Manuela C. Aguirre-Botero, Marek Cyrklaff, Charlotta Funaya, Friedrich Frischknecht

## Abstract

Microtubule inner proteins, MIPs, are microtubule associated proteins that bind to tubulin from the luminal side. MIPs can be found in axonemes to stabilize flagellar beat or within cytoplasmic microtubules. *Plasmodium* spp. are the causative agents of malaria that feature different forms across a complex life cycle with both unique and divergent microtubule-based arrays. Here we investigate the role of four MIPs in a rodent malaria parasite for their role in transmission to and from the mosquito. We show by single and double gene deletions that SPM1 and TrxL1, MIPs associated with the subpellicular microtubules are dispensable for transmission from the vertebrate host to the mosquito and back. In contrast, FAP20 and FAP52, MIPs associated with the axonemes of gametes, are essential for transmission to mosquitoes but only if both genes are deleted. In the absence of both, FAP20 and FAP52 the B-tubule of the axoneme partly detaches from the A-tubule resulting in the deficiency of axonemal beating and hence gamete formation and egress. Our data suggest that a high level of redundancy ensures microtubule stability in the transmissive stages of *Plasmodium*, which is important for parasite transmission.

## Introduction

Microtubule-associated proteins (MAPs) play a vital role in controlling the growth and shrinkage of microtubules, as well as in facilitating the movement of molecular motors along microtubules and securing microtubules in place. Most characterized MAPs bind to the outside of the microtubule or at its ends. Recently, an increasing number of proteins have been found that bind to the microtubule lattice from the inside of the microtubule and have been termed luminal or microtubule inner proteins (MIPs) (*1–15*). Nearly all MIPs have been initially identified in axonemes, including in sperm of metazoans and sea urchins or in flagella of protozoans (*3, 11, 12, 15–18*). Furthermore, several dozen of MIPs have been described in *Chlamydomonas, Tetrahymena* and sperm with mammalian sperm exhibiting the largest number of MIPs discovered so far (*14, 15, 18*). Deletion of some of these MIPs were shown to affect flagellar or ciliar beating pattern (*3, 14, 15*). MIP depletions can result in severe diseases as shown for the flagellar associated protein 52 (FAP52): While *fap52* knockdown leads to severe hydrocephalus in zebrafish, mutations in its human ortholog WDR16 result in human ciliopathies (*14, 19, 20*).

Like other eukaryotes, human infecting protozoan parasites can contain intriguing microtubule arrays (*21, 22*) and feature MIPs within axonemes (*13, 23*) and cytoplasmic microtubules (*1, 21, 24*). In *Plasmodium* spp., the causative agents of malaria in humans and many vertebrates (Figure 1A), cytoplasmic microtubules are closely associated to the membranes of the pellicle, the outer multi-membrane layer of the parasite that surrounds most of the parasite cytoplasm and consists of the parasite plasma membrane and the inner membrane complex (IMC) (Figure 1B). The cytoplasmic microtubules form at the apex of these highly polarized parasites and are closely associated to the IMC by a number of proteins (*25–28*). Due to this association with the IMC, they are called sub-pellicular microtubules (spMTs). spMTs form in different stages of the parasite and seem important for parasite shape. The number of spMTs varies largely between different stages of *Plasmodium* as well as between different *Plasmodium* species (Figure 1A,B): In merozoites, the forms infecting red blood cells, up to 9 spMTs are found in *P. berghei* compared to only 2-4 spMTs in *P. falciparum* with both lacking any detectable MIPs (*21, 29*). Furthermore, also gametocytes differ in spMTs between the different *Plasmodium* species. While *P. falciparum* contain 21 spMTs and appear in a falciform parasite shape, *P. berghei* gametocytes are round and lack an IMC and spMTs (*27*). In contrast there are over 50 spMTs established in ookinetes, the parasite stage entering the mosquito midgut epithelium, and over 11 spMTs are assembled in sporozoites, the forms transmitted by the mosquito. Both forms need to migrate through different tissue barriers, which might require especially stable microtubules. spMTs in ookinetes and sporozoites both feature MIPs arranged in a hemi-spiral (Figure 1C) (*21*). These MIPs were first identified in the related parasite *Toxoplasma gondii* (*T. gondii*) as SPM1 (subpellicular microtubule colocalizing protein 1) and TrxL1 (Thioredoxin-like 1) and shown to fit the luminal densities found in *Plasmodium* (*21, 24, 30, 31*). spMTs are not only found in Apicomplexa, the group that both *Toxoplasma* and *Plasmodium* belong to, but also in the unrelated kinetoplastid parasites *Leishmania* and *Trypanosoma* that lack an IMC. In all these parasites the spMTs are non-dynamic, highly stabilised and cannot be depolymerised by classical methods such as drug- or detergent treatment. We and others thus argued that MIPs in Apicomplexan spMTs might contribute to microtubule stability (*1, 21, 24*). Indeed, parasites lacking MIPs in *T. gondii* show less stable microtubules upon detergent extraction (*24*). Yet, the *T. gondii* parasites lacking MIPs appeared to grow just fine in tissue culture (*24, 32*). However, it is worth noting that in tissue culture *T. gondii* only need to invade cells while sporozoites need to cross barriers such as the basal membrane around salivary glands or the skin during transmission. It remains thus unclear if SPM1 and/or TrxL1 are required for microtubule stability in *Plasmodium* transmissions stages.

**Figure 1:**
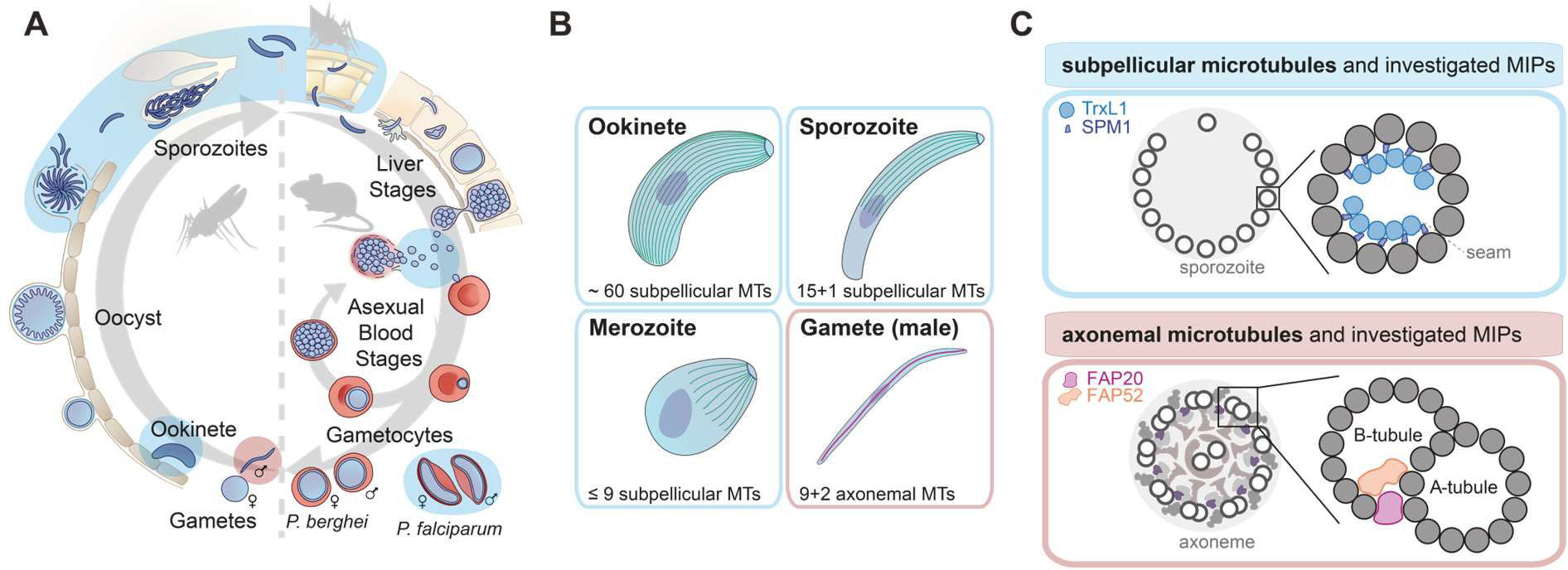
*Plasmodium* relies on different types of microtubules for life cycle progression. **A)** *Plasmodium* life cycle. Forms that possess spMTs are highlighted in light blue, forms that possess axonemal MTs are highlighted in red. Note that *P. berghei* and *P. falciparum* form differently shaped gametocytes with only *P. falciparum* gametocytes containing spMTs. **B**) Schematic of representative *P. berghei* forms containing spMTs (blue box) or axonemal MTs (male gamete). spMTs are shown in green, axonemal MTs in magenta, nucleus in purple, apical polar ring in blue. To allow for better visibility, not all MTs are shown. **C)** Close-up on spMTs with predicted localisation of SPM1 and TrxL1 (based on (*21*)) and axonemal MTs with predicted localisation of FAP20 and FAP52 (based on (*10, 14, 15, 18*)).

*Plasmodium* parasites develop sexual forms, the gametocytes (Figure 1A). Gametocytes that are taken up during a mosquito blood meal rapidly differentiate in the midgut of the insect and exit their red blood host cell (*33*). Male gametocytes undergo three rounds of rapid (<10 min) genome replication via closed mitosis and coordinate the formation of eight axonemes with genome multiplication and separation (*34*). From a bipartite microtubule organizing center (MTOC) that bridges the nuclear envelope, mitotic spindles form within the nucleus, and axonemes form at the cytosolic face of the MTOC (Figure 1B). *Plasmodium* axonemes usually feature a classic 9+2 microtubule array with two central microtubules being surrounded by 9 doublets (Figure 1C). Yet, often axonemes with deviating arrays are formed (*35*). Parasites that cannot form axonemes or form non-motile axonemes are infertile and cannot establish an infection in the mosquito (*36–38*). Whether *Plasmodium* axoneme assembly, stability or function depends on MIPs has not been studied so far.

To investigate the role of MIPs in *Plasmodium*, we generated single and double gene deletion mutants lacking *spm1* and *trxL1* or the flagella associated proteins *fap20* and *fap52*, respectively. We investigated the progression of the clonal parasite lines along the life cycle in the rodent malaria parasite *P. berghei* and found that individual deletions cause no problem for parasite transmission to or from the mosquito. While also the combined deletion of *spm1* and *trxL1* did not hamper transmission to or from mosquitoes, the deletion of both *fap20* and *fap52* led to an almost complete block of transmission. These double mutants formed axonemes which remained non-motile and no gametes emerged. Electron tomography revealed that the lack of motility was caused by the partial detachment of the B-tubule from the A-tubule in axonemes.

## Results

### Deletion of *spm1* and *trxL1* does not affect parasite transmission to and from mosquitoes

To investigate the role of SPM1 and TrxL1 in *P. berghei*, we generated two mutant parasite lines *spm1*(-) and *trxL1*(-) lacking the respective genes and a mutant parasite line lacking both genes *trxL1*(-)*/spm1*(-) (Supplementary Figure 1, 2). Transgenesis and selection of clonal parasite lines in *Plasmodium* spp. is performed in the rapidly multiplying blood stages (*39–41*). After transfection of the knock-out constructs, we readily obtained three clonal lines and all parasite lines grew at normal rates in blood stages (Figure 2A). To test infection of mosquitoes we allowed *Anopheles stephensi* mosquitoes to bite mice infected with either of the mutants or with wild type parasites. 11-12 days later we evaluated the number of oocysts formed in the infected mosquitoes (Figure 2B). All three MIP mutants showed oocysts counts slightly higher than wild type but with comparable midgut infection rates. This suggests that SPM1 and TrxL1 are dispensable for *in vivo* migration into mosquito midguts and establishing of infection. *Plasmodium* sporozoites develop within oocysts in a process depending on microtubules (*42*).

**Figure 2:**
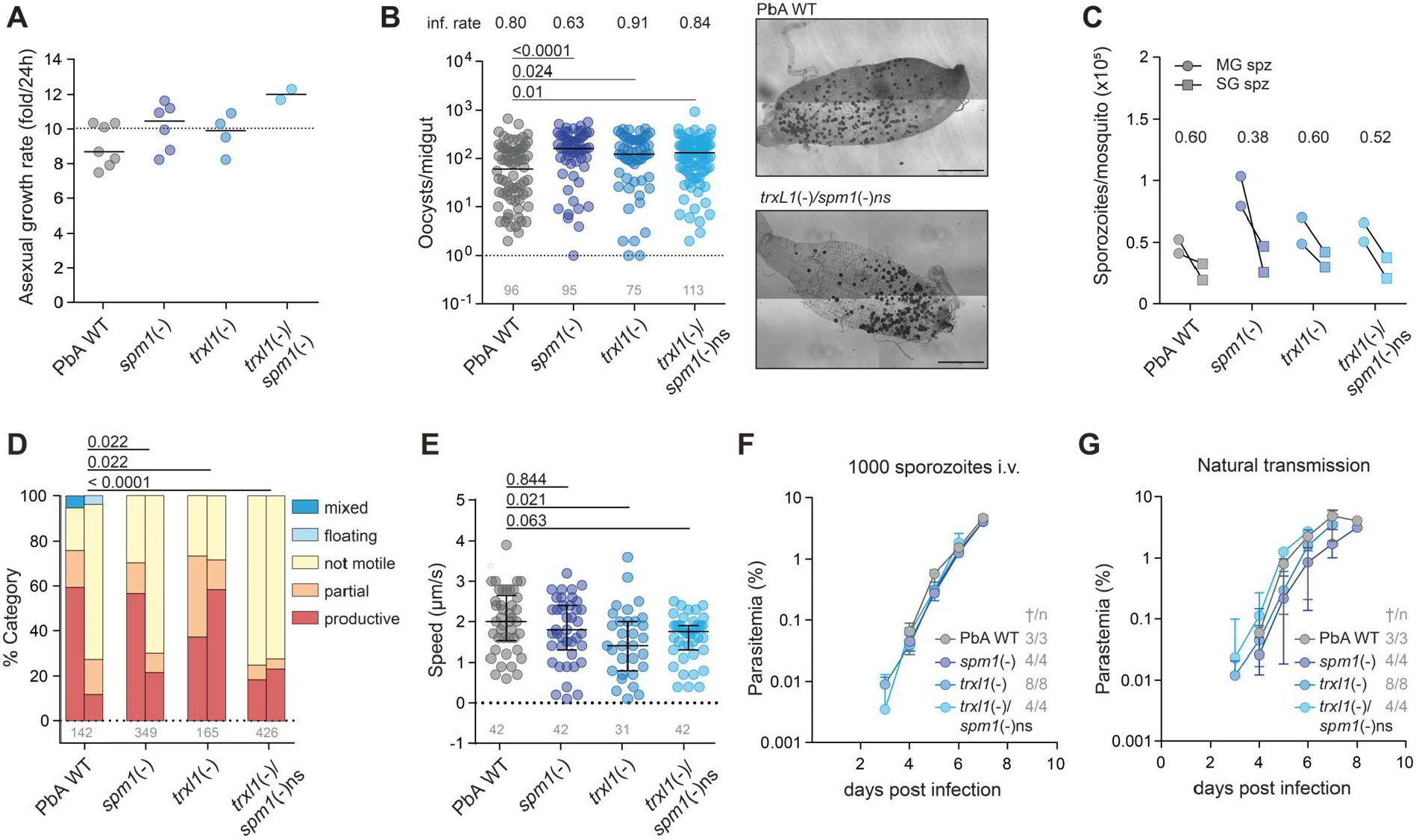
Depletion of *spm1* and *trxL1* does not affect parasite life cycle progression. **A)** Asexual blood stage growth rate as calculated after injecting mice with one iRBC intravenously. PbANKA WT growth rates determined and previously published by our laboratory (*43–45*) are plotted as reference. Line indicates median. **B)** Oocyst counts per midgut. Pooled data from two cage feeds, each dot representing one midgut. Black line indicates median. Mean infection rates are indicated above the graph. Grey numbers indicate total number of midguts analysed. Kruskal-Wallis with Dunn’s multiple comparisons test. Representative image of a wildtype-versus a mutant-infected midgut are shown with oocysts stained in black. Scale: 500 μm. **C)** Salivary gland and midgut sporozoite counts and ratio. Ratios of salivary gland/midgut sporozoites are depicted at the top. **D)** Gliding motility patterns of salivary gland sporozoites. Each bar represents an independent replicate. Statistics: multiple Fisher’s exact tests comparing moving (pooled counts of productive and partial movers) versus non-moving (pooled counts of non-mover, floating and mixed) sporozoites, adjusted for multiple testing according to Bonferroni-Holm. **E)** Speed of productively moving salivary gland sporozoites. Kruskal-Wallis with Dunn’s multiple comparisons test. **F, G)** Parasitemia after infection with F) 1000 sporozoites intravenously or G) natural transmission by bite. Numbers (t/n) indicate blood-stage positive mice vs total mice.

Further, assembling the correct number of microtubules is important for their infectivity of salivary glands and their ability to infect mice (*42*). Investigating the three mutants we found that all formed similar numbers of sporozoites in oocysts and that all could colonize the salivary glands to similar levels (Figure 2C). Sporozoites move at high speeds and thus deletion of genes often affects their gliding motility (*43, 46–48*). To investigate gliding activity, we allowed salivary gland derived sporozoites to settle on a glass substrate and imaged their movement. Quantification of migration patterns and speed revealed a decrease in speed in *trxl1*(-) sporozoites but neither in *spm1*(-) nor in *trxL1*(-)*/spm1*(-)ns ones (Figure 2D, E). To test for the parasite’s transmission capability to mice, we injected either 1000 salivary gland derived sporozoites intravenously into naїve C57Bl/6 mice or let 10 infected mosquitoes bite on a single naїve mouse. We quantified blood stage infection (parasitemia) from day 3 post infection onwards and found no relevant difference between the mutants and wild type parasites (Figure 2F, G). These data together show that the MIPs SPM1 and TrxL1 are dispensable for both transmission from mice to mosquitoes and from mosquitoes to mice.

### Deletion of both *fap20* and *fap52* abrogates transmission to mosquitoes

Most MIPs so far were found in axonemes with over 60 MIPs being revealed in sperm axonemes (*7, 18*). To investigate if there are MIPs in *Plasmodium* axonemes we searched for orthologs to known MIPs from other organisms using PlasmoDB. Blasting known axoneme associated MIPs), we identified orthologues to the two flagellar associated proteins FAP20 and FAP52 ((Supplementary Figure 3, Figure 1C). FAP20 is a highly conserved protein across ciliated organism that attaches with an 8 nm periodicity along the microtubule at the junction of the A and B-tubule (*5*). Deletion of *fap20* in *Chlamydomonas* led to less stable axonemes and an abnormal waveform of the beating cilia resulting in a motility defect (*5, 10, 49*). The identified *P. berghei* and *P. falciparum* FAP20 orthologues are well conserved on sequence and structure level to FAP20 proteins of representative model organisms, but notably carry an extensive, C-terminal loop whose structure is not well predicted (Supplementary figure 3 A, C). FAP52 is a luminal protein of 66 kDa associated with a 16 nm periodicity. Deletion of f*ap52* in *Chlamodymonas* showed no defect in motility (*14*). Interestingly, deletion of *fap52* together with *fap20* in *Chlamydomonas* lead to a complete loss of motility and a detachment of the B-tubule from the A-tubule (*14*). *P. berghei* and *P. falciparum* FAP52 orthologues were less well preserved on a sequence level, but their structure, consisting of two WD40 domains, was highly similar to that of human or *Chlamydomonas* FAP52 (Supplementary figure 3 B, D).

To probe for a potential function of FAP20 and FAP52 in *Plasmodium*, we generated single *fap20*(-) and *fap52*(-) and double *fap20/52*(-) gene deletion mutants in *P. berghei* (Supplementary Figure 1, 4) in the gametocyte reporter line Pb820 WT, which expresses cytosolic GFP in male gametocytes and cytosolic RFP in female gametocytes (*50*). After transfection of the knock-out constructs and limited dilution we readily obtained three clonal lines which all grew at normal rates in blood stages and developed gametocytes (Figure 3A, B) To investigate for a potential role of the proteins in axoneme formation or function we activated gametocyte containing blood and investigated the rate of gamete formation (exflagellation) and ookinete formation. We observed that the single gene deletion mutants could readily exflagellate from the red blood cell and form ookinetes with slightly less efficacy of ookinete formation for the *fap52*(-) line (Figure 3C, D). Strikingly, however, the double mutant failed to exflagellate and only formed very few ookinetes (Figure 3C, D. To test for the capacity of the mutants to infect mosquitoes, we allowed mosquitoes to bite on infected mice and scored the numbers of oocysts. Both single mutant *fap* lines infected the midgut and formed oocysts to an extend comparable to wildtype. In comparison, the double knockout mutant had a much-reduced infection rate and barely formed oocysts (Figure 3E). The few oocysts that formed however showed normal oocyst sizes, indicating normal development (Figure 3F). Sporozoites developed in both single mutants inside the oocyst and could infect salivary glands but showed an overall lower sporozoite count (Figure 3G). Despite the slightly lower parasite load in the salivary glands, both single mutant lines were able to transmit and infect mice upon natural transmission by mosquito bite indistinguishable from wildtype parasites (Figure 3H). These data show that single depletion of either *fap20* or *fap52* does not impact mosquito infection or transmission from mosquito to mice. However, depletion of both *faps* affects male gamete differentiation that results in drastically reduced ookinete conversion rates and an almost abolished midgut infection.

**Figure 3:**
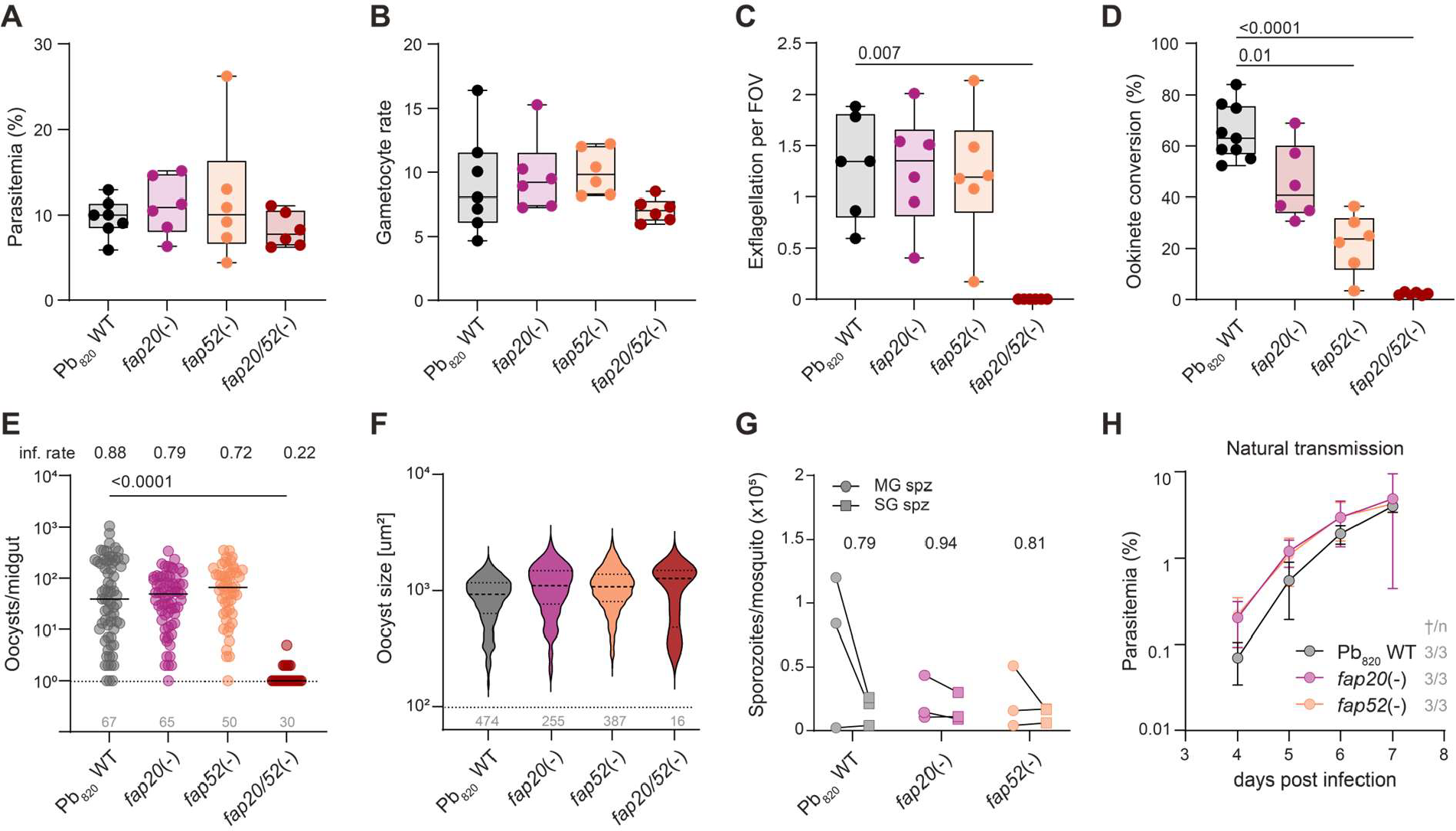
Depletion of both *fap20* and *fap52* prevents productive mosquito infection. **A, B)** A) Parasitemia and B) gametocyte rate 3 days after infecting mice with two Mio iRBC intravenously of the indicated parasite lines. **C)** Exflagellation per field of view (FOV) of the indicated parasite lines. **D**) Ookinete conversion rate in % of all red fluorescent parasites (i.e. females, gametocytes, ookinetes). **E)** Oocyst counts per midgut. Pooled data from three cage feeds, each dot representing one midgut. Black line indicates median. Mean infection rates are indicated above the graph. Grey numbers indicate total number of midguts analysed. **F)** Oocyst size. Pooled data from three cage feeds. Grey numbers indicate total number of oocysts analysed. **G**) Salivary gland and midgut sporozoite counts and ratio. Ratios of salivary gland/midgut sporozoites are depicted at the top **H)** Parasitemia after infection by natural transmission. Numbers (t/n) indicate blood-stage positive mice vs total mice. **C, D, E)** Statistics: Kruskal-Wallis with Dunn’s multiple comparisons test.

### The B-tubule detaches from the A-tubule in mutants lacking both *fap20* and *fap52*

To identify the cause for the lower exflagellation rate and subsequent impaired midgut infection in the double mutant, we investigated organization of axonemal microtubules in activated gametocytes. For this, we fixed gametocytes 15 minutes post activation, prepared them for transmission electron microscopy and took electron micrographs from thick 200 nm sections (Supplementary figure 5). Blinded analysis of these images demonstrated the presence of axonemes in male gametocytes, but we noticed in some microtubule doublets of *fap20/52*(-) a small gap between the protofilaments that link the B-tubule to the A-tubule (Figure 4A). This gap was only rarely seen in the single mutants and not in the wild type (Figure 4B). As the gap was only apparent in some of the *fap20*/*52*(-) axonemal doublets we hypothesised that either some doublets have no such gap, while others do, or that all doublets show a gap at some sections, but not along the entire axoneme. To distinguish between these scenarios, we employed electron tomography of 200 nm thick serial sections from the double mutant (Supplementary figure 6). To best view the axonemes we projected 20 virtual sections, which revealed the axonemes in enough clarity to distinguish doublets and singlets (Figure 4C, Supplementary figure 7A-C). Still, in only a variable fraction of sections (from 3 % to 61 %) were we able to clearly distinguish between attached or detached B-tubules, while the majority showed too little contrast to confidently distinguish these two states (Figure 4E, F, Supplementary figure 7D-F, Table 1). Yet, despite these challenges, we could follow the attachment state of the B-tubule along all doublets (Figure 4D, Supplementary Figure 8, Table 1). This showed that for most of the 90 examined microtubule doublets (from 10 examined axonemes from three different gametocytes) the B-tubule could be detached from the A-tubule for as little as 20 nm to at least 500 nm. Similarly, the B-tubule was found to be attached for as few as 20 nm between two states of detachment or for as long as at least 500 nm (Figure 4D, Supplementary Figure 8). Axonemes in *Plasmodium* gametes do not always show a 9+2 pattern and we also observed a number of different arrangements in the double mutants (Supplementary figure 9). In one instance we found a complete detachment of a B-tubule from the A-tubule (inner and outer junction detached) prior to re-attachment and in another one we could follow a splaying (or disassembling) axoneme tip (Supplementary figure 9 C,D). Together these data show that in the *fap20/fap52*(-) mutant B-tubules partially detach from the A-tubules leading to non-motile axonemes and a near complete block in exflagellation, which essentially stops parasite transmission from the vertebrate to the mosquito. Given that such a defect was not observed in the single *fap* depletion mutants, these results demonstrate that FAP20 and FAP52 can compensate for each other to form the link between A- and B-tubule (Figure 4G).

**Figure 4:**
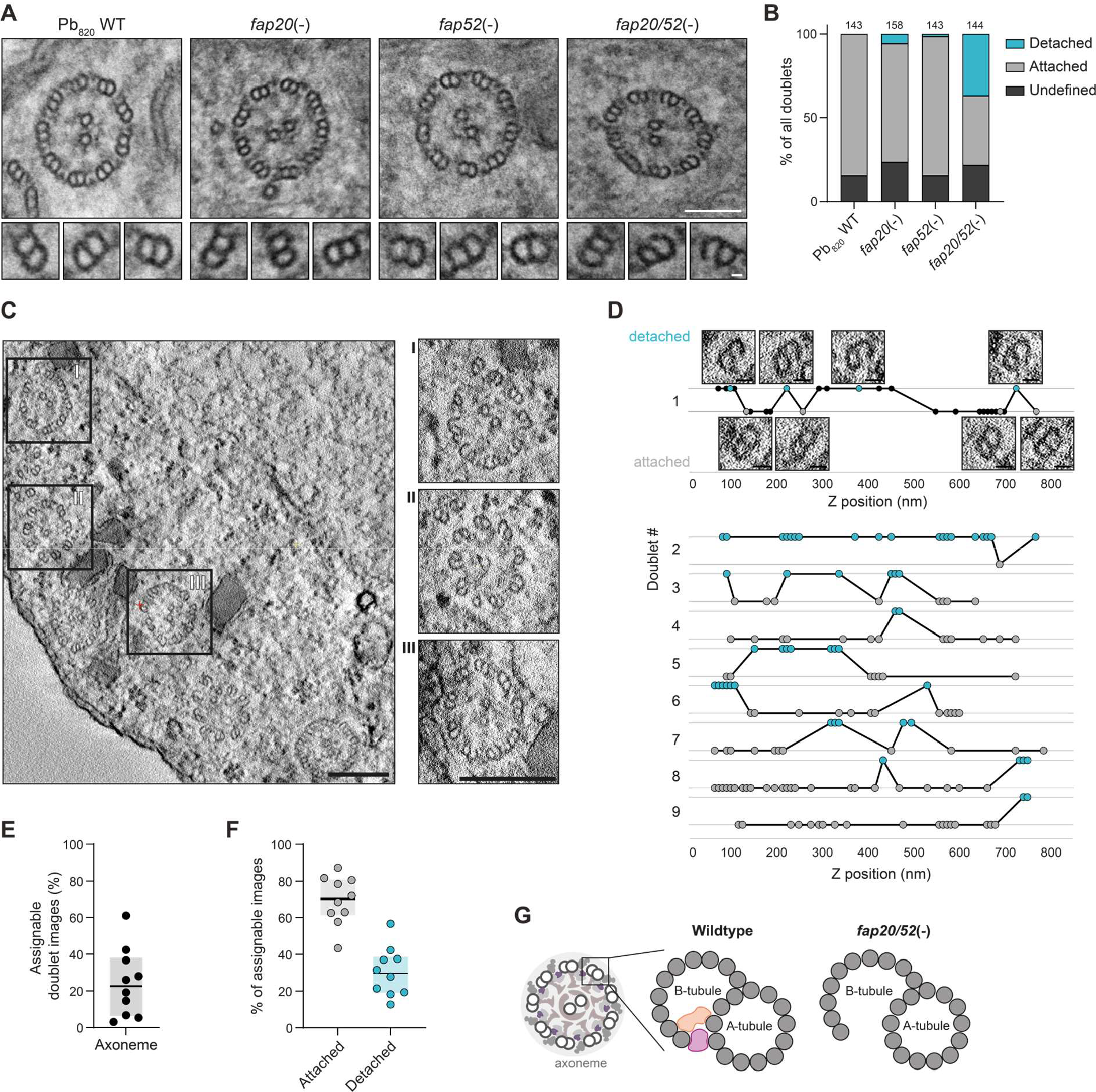
Depletion of both *fap20* and *fap52* leads to a detachment of the B-tubule from the A-tubule. **A)** Example TEM sections of individual axonemes (upper row, scale bar: 100 nm) and doublets (lower row, scale bar: 10 nm) in gametocytes fixed 12 min after activation (mpa). **B)** Frequency of doublet states in TEM sections. **C)** Example tomogram section of *fap20/52*(-) gametocyte 12 mpa. with close-up of investigated axonemes I-III. Scale bar: 200 nm. Sum projection of 20 z-layers. **D)** Doublet states of doublets 1 to 9 of axoneme I across z position. Each dot represents an assignable doubled image. Turquoise, detached, grey, attached. First row, example images corresponding to coloured dots are shown, black, data points without example picture. Scale bar: 20 nm. **E)** Fraction of assignable doublet images. Each dot represents one axoneme. **F)** Fraction of attached or detached doublets of all assignable images. Each dot represents one axoneme. **G)** Scheme of proposed doublet state in wildtype and *fap20/52*(-) parasites.

**Table 1:**
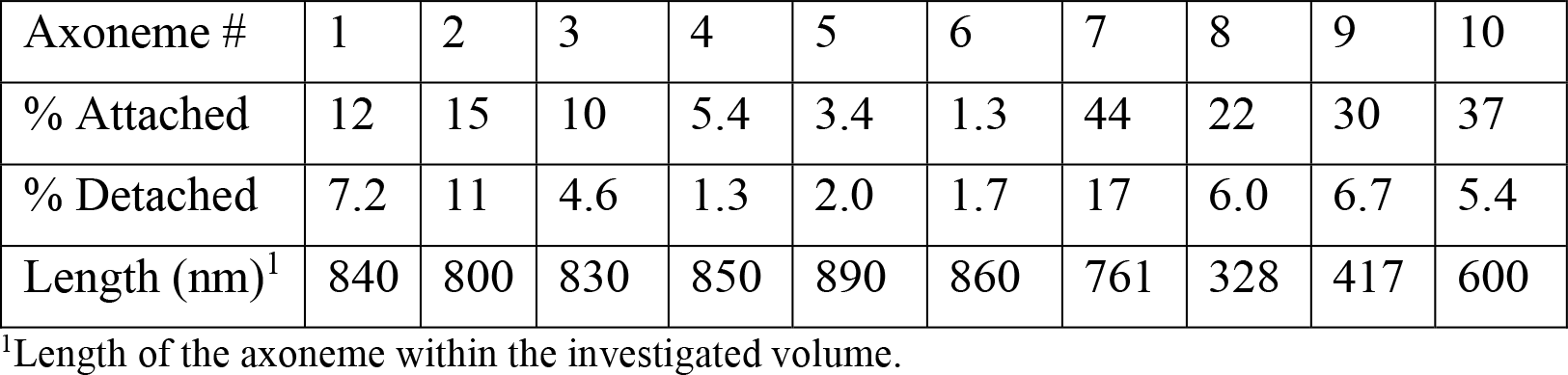
Doublet states as percentage of all doublets along individual axonemes.

## Discussion

Microtubules are key scaffolding proteins to give parasites their shape and mediate the right balance between cellular elasticity and rigidity. It is widely assumed that microtubule inner proteins (MIPs) contribute to microtubule stability and that deletion or loss of them will cause weakened microtubules. As microtubules are key for the formation and shape of most of the different extracellular forms of the malaria parasite, we here investigated four *Plasmodium* MIPs for their potential function in life cycle progression by generating single and double deletion lines. Two of the MIPs, SPM1 and TrxL1, were identified in the subpellicular microtubules of ookinetes and sporozoites and two, FAP20 and FAP52, in the axonemes of activated male gametocytes/gametes.

Deletion of *spm1* and *trxL1* individually or in combination did not affect the capacity of ookinetes or sporozoites to migrate and establish infection. This suggests that additional factors are likely important for ensuring microtubule stability. These could be the stabilizing effect of acetylation of the luminally exposed lysine 40, which was indeed suggested to be important for microtubule stability in *T. gondii* (*51*). Also, differences in the contacts between the tubulin dimers making up the protofilaments of microtubules could lead to overall more stable microtubules. The fact that the double deletion of *spm1* and *trxL1* is not showing a more prominent phenotypic difference to wild type than the single *spm1* deletion is not surprising as TrxL1 was shown to depend on SPM1 for binding to microtubules (*24*). Initial work in *T. gondii* identified three MIPs in spMTs: SPM1, TrxL1 and TrxL2 (*24*). Of these only the first two are conserved in *Plasmodium* spp. Yet, also in *T. gondii* the deletion of all MIPs showed no phenotypic difference to wild type parasites during parasite culture in human fibroblasts. A difference was only found after chemical extraction of the microtubule cytoskeleton, a somewhat non-physiological condition. We hoped that deletion of the MIPs in *P. berghei* would reveal a difference considering the strenuous migrations these parasites need to undergo. Ookinetes move only across a single cell layer and have many microtubules. They might hence be more accommodating for minor permutation than the sporozoites. Sporozoites need to cross the basal lamina of the salivary gland cell, a formidable barrier that requires considerable force (*43, 52*). Furthermore, the sporozoites need to migrate in the skin, during which the parasites can take sharp turns necessitating a strong subpellicular MT cage that provides high degree of cell stability (*48, 53*). Nevertheless, we could not detect a difference in establishing an infection during transmission from mosquito to mouse, indicating that this cell stability is not only conferred by these two MIPs (Figure 2 F, G).

The individual deletions of *fap20* and *fap52* also showed no remarkable differences to the wild type in terms of transmission to mosquitoes despite lower sporozoite counts in both the oocyst and the salivary glands. This highlights, that even a much-reduced sporozoite load in the salivary gland is sufficient to induce infection in mice. However, the deletion of both *fap20* and *fap52* nearly completely abrogated transmission as evidenced by a more than 90 % decrease in the numbers of oocysts that could establish in mosquito guts (Figure E). This finding shows that there is an important degree of redundancy in the MIP system. Hence axonemal microtubule stability must be of great importance to the parasite, and it might well be that our assays cannot distinguish subtle differences between the single knockout lines that could be important in a natural setting. Of note, deletion of only *fap52* did reduce ookinete conversion rate. While in laboratory settings, this reduction did not translate into a reduced mosquito infection rate, it is likely that under natural transmission settings, a reduced ookinete conversion rate does affect parasite life cycle progression. Several dozen MIPs have been shown to bind to the lumen of axonemal MTs in other organisms (*10*). Hence it is surprising, that compared to a total of 42 MIPs in *Tetrahymena* or 33 MIPs in *Chlamydomonas* axonemes (*10, 15*), we have so far identified only two axonemal MIPs in *Plasmodium*.

We showed that the combined depletion of *spm1* and *trxL1* did neither affect parasite development nor transmission, it still remains to be elucidated whether there are other MIPs that contribute to the high degree of spMT stability. While computational modelling of SPM1 and TrxL1 enabled to fit in the two proteins into two half crescents similar to the ones shown in *T. gondii* (*21, 24*), the lack of an TrxL2 orthologue in *Plasmodium* suggest another protein to fill into the space between the two half crescents (*32*). Besides MIPs binding to the microtubule lumen also MAPs binding to the microtubule surface should be considered, as they add to the overall spMT stability. Indeed, recent work showed that depletion of three proteins that bind to the cortical MTs in *T. gondii* resulted in much shorter and destabilized MTs (*54*). Even though this defect resulted in lower parasite speed, parasites replicated still normally highlighting the extreme robustness of MT stability in apicomplexan parasites.

In conclusion, we show that microtubule inner proteins are important for transmission from vertebrate host to mosquitoes. While spMTs showed to be insensitive to double depletion of two MIPs, axonemal MT in *Plasmodium* are dependent on FAP20 and FAP52 for both gametocyte maturation and midgut infection to occur.

## Materials and Methods

### Ethics statement

Animal experiments were performed according to Federation of European Laboratory Animal Science Associations (FELASA) and Society of Laboratory Animal Science (GV-SOLAS) guidelines and approved by the responsible German authorities (Regierungspräsidium Karlsruhe). Mice were obtained from Janvier or Charles River Laboratories and kept in the dedicated animal facility of Heidelberg University according to current guidelines (3 mice per cage, *ad libitum* food and water, environmentally enriched cages).

### Animals and parasites

Female 4–6-week-old Swiss CD1 mice from Janvier were used to generate mutant *P. berghei* parasite lines based on the parental *P. berghei* ANKA strain (*55*) or the *P. berghei* 820 line (generous gift from A. Waters, University of Glasgow, (*50*)), to propagate parasites and to generate gametocytes or infect *Anopheles stephensi* mosquitoes for sporozoite production. For transmission experiments to determine sporozoite infectivity, 4-6-week-old C57/BL6 mice from Charles River laboratories were used. *Anopheles stephensi* mosquitoes were reared and maintained according to standard procedures.

### Cloning of plasmid construct for parasite transfection

For gene deletion of *Pb*SPM1 (PBANKA_0810700), *Pb*TrxL1 (PBANKA_0820200), *Pb*FAP20 (PBANKA_1108000) and *Pb*FAP52 (PBANKA_1361500), the respective transfection vectors were obtained from PlasmoGEM (*56, 57*). These PlasmoGEM vectors contain resistance cassettes allowing both for positive and negative selection to recycle the resistance marker (*58*). Prior to transfection, 3-5 μg of each vector were linearized using NotI followed by ethanol precipitation.

### Generation of *P. berghei* parasite lines

*P. berghei* single and double-knockout mutants were generated via double homologous crossover recombination (Supplementary Figure 1A,B). For this, the linearized vectors were transfected into the parental *P. berghei* strain (PbANKA wildtype for SPM1 and TrxL1, and Pb820 wildtype for FAP20 and FAP50) using standard protocols (*39*). Transgenic parasites were selected for via pyrimethamine (0.07 mg/mL, positive selection) administered via the mice’s drinking water. To obtain clonal lines from the mixed parasite populations after transfection, a limiting dilution was performed. For this, a single-blood stage parasite was injected into each of 4 to 8 Swiss mice intravenously. Once mice reached 1-3 % parasitemia, blood was collected via cardiac puncture from anesthetized mice (120 mg/kg ketamine, 16 mg/kg xylazine, intraperitoneally). Cryostabilates were stored in liquid nitrogen. For genotyping, parasite-infected blood was lysed in 0.093 % saponin, and parasites were resuspended in 200 μl PBS. Genomic DNA was isolated using a blood and tissue kit (Qiagen Ltd.) according to manufacturer’s protocol. All generated isogenic parasite lines were analyzed via genotyping PCR, with all primers listed in supplementary Table 1. Parasite lines with the correct genotype were assumed to be isogenic.

In order to be able to reuse pyrimethamine as a selective (positive) pressure in subsequent mutant generations, 5-fluorocytosine (5-FC) selection (negative) was performed (Supplementary Figure 1A,B). The yeast enzyme cytosine deaminase and uridyl phosphoribosyl transferase (*yFCU*) gene in the transfection constructs allows for negative selection and hence the loss of the selection cassette as it metabolizes the prodrug 5-FC into the toxic 5-fluorouracil (5-FU)(*59*). Only those parasites that recycled the selection cassette and hence then lack *yFCU* do not convert 5-FC into 5-FU and will survive. Here, *trxL1*(-) and *fap20*(-) parasites that recycled the selection cassette were negatively selected for via 5-FC (1 mg/ml) administered via the mice’s drinking water and isogenic clones were isolated by limited dilution and verified via genotyping PCR accordingly. After obtaining marker-free clones, the second knockout (*spm1* or *fap52*) was generated in these lines as described above.

### Determination of asexual blood stage growth and gametocyte rate

For *spm1*(-), *trxl1*(-) and *spm1*(-)*/trxl1*(-)ns parasite lines, asexual growth rates were determined from all isogenic lines obtained after limiting dilution. Parasitemia on day 8 after injection of a single parasite was used to calculate the growth rate as described before. (*60, 61*). Growth rate calculated for the double KO mutant (Figure 2A) represents the calculated one from the donor line *trxL1*(-)/*spm1*(-) while characterization was carried out with the negatively selected *trxL1*(-)/*spm1*(-)ns clone. For *fap20*(-), *fap52*(-), and *fap20/52*(-) parasite lines, asexual growth was obtained by determining parasitemia from giemsa-stained blood smears 3 days after injection of phenylhydrazine-pretreated mice (see below) with 2*10^6^ iRBC i.v. From the same slides, gametocytemia was determined and gametocyte rate was calculated as the proportion of parasites that were gametocytes over all parasites.

### Analysis of exflagellation rate

Mice were pre-treated with 200 μl phenylhydrazine (6 mg/ml) to increase reticulocytemia and therefore gametocytemia two to three days before intravenous infection with 2*10^6^ iRBC. Three days post infection, exflagellation was determined. For this, one drop of tail blood was mixed with 5 μl ookinete medium (RPMI supplemented with 20 % (v/v) FCS, 50 μg/ml hypoxanthine, and 100 μM xanthurenic acid, adjusted to pH 7.8 – 8.0), placed onto a glass slide and tightly covered with a cover glass. Slides were incubated at 21 °C for 13 min and then imaged immediately at a Zeiss Axiostar microscope at 40x magnification and with a phase contrast ring. Exflagellations and fields of view were counted for 2.5 min in areas with roughly comparable red blood cell densities. Exflagellation rates were calculated as mean of three technical replicates for each infected mouse.

### Ookinete conversion rate and motility

Mice infected for exflagellation assays (see above) were subsequently bled via cardiac puncture and 500 μl blood was transferred to 10 ml ookinete medium. Parasites were incubated at 19 °C for 20-24 h. Ookinetes were then enriched over a 63 % Nycodenz (Serumwerk Bernburg AG, Ref 18003, Bernburg, Germany) cushion as described before (*62*) the pellet was resuspended in 10 μl ookinete medium, and 2-3 μl of the suspension were placed onto a glass slide, tightly covered with a cover glass and imaged at an epifluorescence microscope (Zeiss Axiovert 200M, 25x objective). For ookinete conversion rates, images were taken from randomly selected areas. Ookinete conversion rate was calculated as the number of ookinetes divided by the total number of red fluorescent cells (females, zygotes and ookinetes, based on their red fluorescence in the Pb820 reporter line). For ookinete motility, time courses were taken in the DIC and RFP channel at a frame rate of 1 frame/20 seconds for 15 min/movie and ookinete speed was determined using the FIJI plugin Trackmate 6.0.3 plugin (*63*).

### Mosquito infection

Two naїve Swiss mice were infected with 20*10^6^ iRBC i.p. Three days later, presence of male gametocytes was assessed by determining exflagellation as described above. If a sufficient amount of exflagellation centers were observed, mice were anaesthetised with 100 mg/kg ketamine and 3 mg/ml xylazine i.p. and exposed to approximately 200 female *Anopheles stephensi* mosquitoes. Mosquitoes were allowed to feed for 20 min to 40 min. Infected mosquitoes were kept at 21 °C and 80 % humidity and provided with both sugar and salt pads.

### Analysis of oocyst development by mercurochrome staining

Midguts of infected female mosquitoes were dissected in 100 μl PBS on ice as technical replicates on day 11 and day 12 post infection. Isolated midguts were permeabilized in 1 % Nonidet P40 (Applichem, #A1694, 0250) for 20 min at room temperature, followed by staining with 0.1 % mercurochrome in PBS (NF XII, Sigma-Aldrich) for 30 to 90 min. Midguts were washed multiple times with PBS, placed onto a glass slide and covered with a coverslip. Midguts were imaged at an epifluorescence microscope (Zeiss Axiovert 200M) at 10x magnification (NA 0.5, air) (for infection rate and oocyst count) and at 25x magnification (for oocyst size) with a green filter (38 HE Green Fluorescent Prot). Oocyst size was determined only for day 12 oocysts and measured in FIJI (Vers. 2.9.0).

### Sporozoite isolation, counting and motility assay

Midgut and salivary glands were dissected in PBS as technical replicates between day 18 to day 22 post blood-meal of two independent mosquito cage feeds. A minimum of 15 female mosquitoes were dissected per replicate. Isolated organs were crushed with a pestle for 1 min to release sporozoites following counting using a Neubauer counting chamber.

To perform gliding assays, isolated salivary gland sporozoites (in 50 μl RPMI) were purified via an Accudenz-based gradient centrifugation. For this, the salivary gland sporozoite suspension was topped up to 1 ml with RPM and then carefully underlaid with 3 ml of 17 % Accudenz (in dH_2_O, Accurate Chemical & Scientific Corp., Westbury, NY, USA). Centrifugation for 20 min at 2800 rpm and at room temperature without break separated sporozoites from cell debris. The sporozoite containing interphase was taken-off carefully (total of 1.4 ml) to directly pellet sporozoites for 3 min at 13’000 rpm at room-temperature. Sporozoites were activated in 3 % BSA/RPMI (Carl Roth GmbH + Co. KG, Karlsruhe, Germany) and transferred into a 96-optical well-plate (Thermo Fisher Scientific, Waltham, MA, USA). The plate was spun for 3 min at 1’000 rpm at room temperature. Sporozoites were imaged at an epifluorescence microscope (Zeiss Axiovert 200M) with a 25x objective (NA 0.8, oil) in DIC at a frame rate of 3s for 100 cycles until latest 1 hour post activation. Sporozoite gliding behaviour was then analysed single-blinded using FIJI (Vers. 2.14.0) and categorized into four different patterns as described before (*64*): productive: persistently moving sporozoites for at least 50 frames with less than 10 frames of stopping, partial: sporozoites that moved more than one parasite length but less than 50 frames, attached: sporozoites that stayed attached at the surface and moved less than one parasite length, floating: sporozoites that detach without contact to the substrate. Productive movers were then further analysed for their speed by back calculating from the number of circles moved/ number of time frames and the circle diameter.

### Transmission to mice

For natural transmission via mosquito bite, ten female mosquitoes (day 20 post infectious blood meal), were transferred into cups the day before the experiment. Mosquitoes were starved for at least 3 hours prior to feed on the day of the experiment. Per replicate, four six-weeks old naїve C57/BL6 mice were anaesthetised with 120 mg/kg ketamine and 16 mg/ml xylazine via intraperitoneal injection and one mouse each was placed onto a mosquito-containing cup. Mosquitoes were allowed to feed for at least 15 min up to 30 min a minimum number of six mosquitos to have fed. Afterwards, the salivary glands from all blood-fed mosquitos were harvested to check for the presence of sporozoites. For intravenous injection of sporozoites, salivary glands were dissected as previously described and 1’000 sporozoites (in 100 μl PBS) each were injected into a lateral teil vein of four naїve C57/BL6 mouse. Blood smears from a drop of tail blood were collected daily from day 3 post infection on. Once mice reached a parasitemia of 2-3 %, they were sacrificed via cervical dislocation. Mice that did not become positive within 20 days post infection were considered negative.

### Gametocyte fixation and electron microscopy

Mice with more than 3 % parasitemia were bled, blood was collected into 5 ml prewarmed medium and gametocytes were purified over a 49 % nycodenz cushion (25 min, 1500 rpm, no brake). All steps were kept at 37 °C to prevent premature activation. Gametocytes were collected from the interphase and washed once in 8 ml warm medium. To activate gametocytes, they were resuspended in 500 μl ookinete medium and incubated for 15 min at 19 °C. Cells were pelleted and chemically fixed in 4 % paraformaldehyde/ 1 % glutaraldehyde in 100 mM PHEM buffer (240 mM PIPES, 100 mM HEPES, 40 mM EGTA, 8 mM MgSO_4_) at 4 °C overnight. Samples were washed thrice in 100 mM PHEM buffer by incubating samples for 40 seconds in the microwave (BioWave Pro+, PELCO, Fresno, CA, USA) and pelleted for 2 min spin at 2000 rpm). A secondary fixation was done once in 1 % osmium tetroxide (in 100 mM PHEM buffer) for 14 min at 50 °C. Osmium tetroxide was removed washing sample twice in distilled water at RT (5 min spin at 2000 rpm) and then contrasted with 1 % uranyl acetate (in ddH_2_O) at 20 Hg vacuum for 14 min at 50 °C. Samples were washed twice with ddH_2_O for 10 min and then dehydrated for 40 seconds at 50 °C in the microwave in increasing concentrations of aqueous acetone (30 %, 50 %, 70 %, and 90 %) and twice in 100 % dried acetone for 40 seconds at 50 °C in the microwave. Samples were adapted to Spurr’s solution (23.6 % epoxycyclohexyl-methyl-3,4-epoxycyclohexylcarboxylate [ERL], 14.2 % ERL-4206 plasticizer, 61.3 % nonenylsuccinic anhy-dride, 0.9 % dimethylethanolamine) by incubating in increasing concentrations of Spurr resin (25 %, 50 %, 75 % in acetone) for 40 seconds each in the microwave (2 min spin at 2000 rpm) and finally embedded in 100 % Spurr resin at 60 °C for at least 24 hours. Embedded gametocytes were sectioned using an ultramicrotome (LEICA EM UC7, Leica Microsystems GmbH) and first 200-nm-thick single sections were acquired followed by 200 nm-thick continuous sections. Sections were imaged on a transmission electron microscope at 200 kV (FEI Tecnai F20, FEI/Thermo Fisher Scientific, Hillsboro, OR, USA) using a Eagle 4k x 4k CCD camera (FEI Eindhoven, Netherlands). Following image acquisition described before(*65*), images were 2x binned to reduce file size. For tomography, bidirectional single axis tilt-series were taken from -60 ° to +60 ° with 2 ° increments at 25000x magnification resulting in a pixel size of 0.4459 nm. Single images were analysed with FIJI in a blinded fashion (Version 1.53q), while tilt-series were reconstructed and analysed using IMOD (*66*). Doublet MTs were manually categorized into attached/detached/unassignable.

## Data availability

All data have been made available in the manuscript.

## Acknowledgements/funding

We thank Steffi Gold for help in EM preparations and Markus Ganter for comments on the manuscript. We thank Katharina Röver for her help in parasitemia determination and performing genotyping PCRs. We thank Miriam Reinig and all students helping with the mosquito rearing. This project was funded by grants from the Deutsche Forschungsgemeinschaft (DFG, German Research Foundation): SPP 2225 “Exit pathways of intracellular pathogens” (FR2140/12-1), and DFG FR2140/10-1. MS received a visiting fellowship from the Carl Duisberg foundation. AB is a member of the Heidelberg Biosciences International Graduate School (HBIGS). LH is a member of the Molecular Biotechnology Master program, LPD is a member of the Heidelberg Biosciences Infectious Disease Master Program, MA was a member of the Heidelberg Bioscience Molecular and Cellular Biology Master Program. We acknowledge the microscopy support from the Infectious Diseases Imaging Platform (IDIP) at the Center for Integrative Infectious Disease Research and are grateful for the generous use of the microscopes at the Electron Microscopy Core Facility (EMCF) of Heidelberg University. The Plasmodium database PlasmoDB facilitated this work.

The funders had no role in study design, data collection, and interpretation, or the decision to submit the work for publication.

## Authors contributions

Conceptualization: FH, AMB, FF

Methodology: FH, AMB, FF

Investigation: FH, AMB, LPD, LH, FN, MS, MCAB, MC, CF

Writing original Manuscript: FH, AMB, FF

Review & Editing: FH, AMB, FF

Funding acquisition: FF

Resources: FF

Project administration: FF

Supervision: FH, AMB, FF

All authors read and approved the manuscript.

