## Supplementary Figures and Table for "Microtubule inner proteins of *Plasmodium* are essential for transmission of malaria parasites"

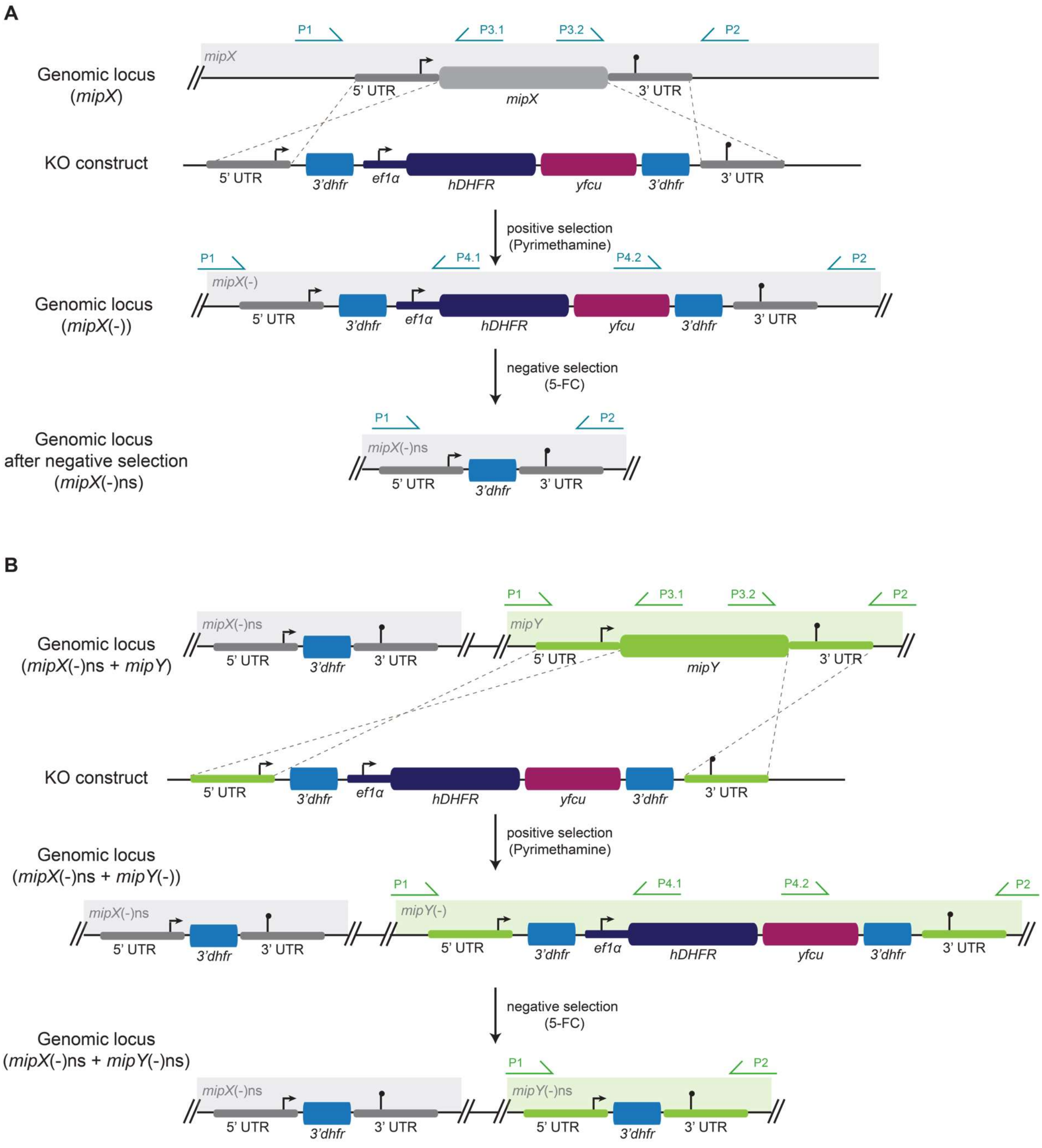


**Supplementary Figure 1: Genetic strategy of generating double knockouts.** The original genomic locus is shown on top with dotted lines indicating the double crossover events leading to insertion of the KO construct. This yields the intermediate parasite line that upon negative selection loops out the resistance cassette by excision across the two flanking *3'dhfr* regions (blue). **A**) Scheme of generation of single knockout including negative selection. Primers used for genotyping PCR are indicated. **B**) Scheme of generating second knockout. Primers used for genotyping PCR are indicated. A, B) ef1α, human elongation factor-1 alpha, hDHFR, human dihydrofolate reductase, KO, knockout, mipX, microtuble inner protein X, ns, negative selected, UTR, untranslated region, yfcu, yeast enzyme cytosine deaminase and uridyl phosphoribosyl transferase. Schemes are not drawn to scale, primer positions are approximate.

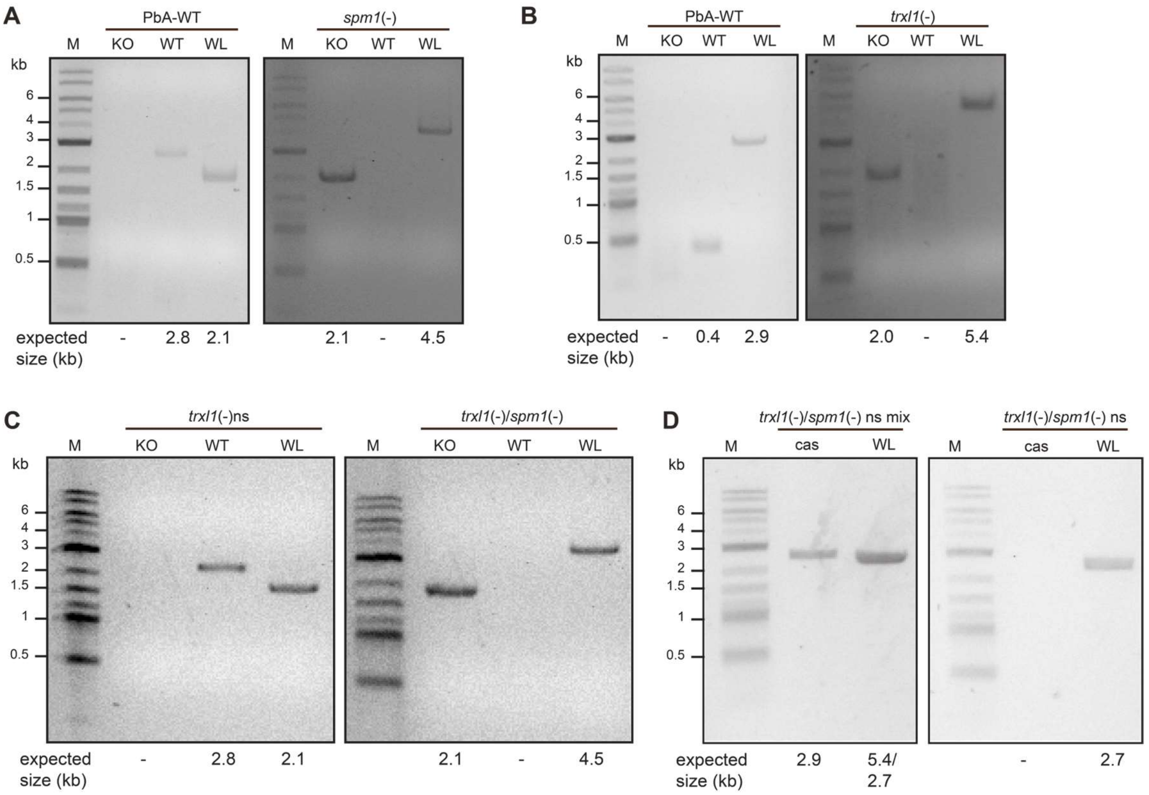


**Supplementary figure 2: Generation of *spm1* and *trxl1* single and double knockout parasite lines.** Genotyping PCRs of **A**) *spm1*(-), **B**) *trxl1*(-), **C**) *trxl1*(-)/*spm1*(-) and **D**) *trxl1*(-)/spm1(-) ns. Primers are indicated in supplementary figure 1 and supplementary table 1. Kb, kilobase, ns, negatively selected

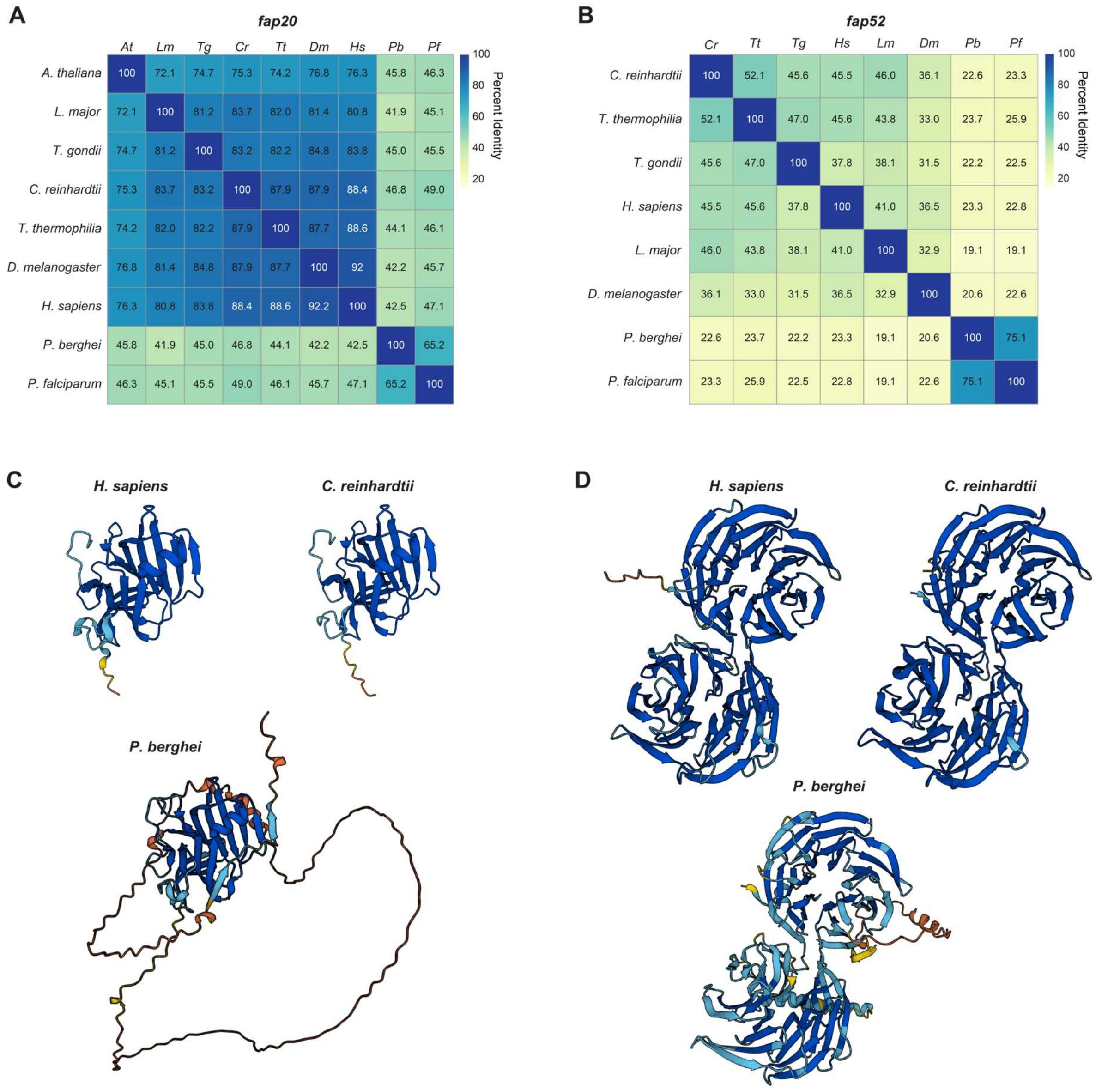


**Supplementary figure 3: Identification of FAPs by homology search. A, B)** Percent identity matrix of A) *fap20* and B) *fap52* genes in *Plasmodium* and selected model species. **C, D)** Alphafold2-predicted structures of A) FAP20 and B) FAP52 in *H. sapiens*, *C. reinhardtii*, and *P. berghei* (*1*, *2*). Note the large extension for *P. berghei* FAP20.

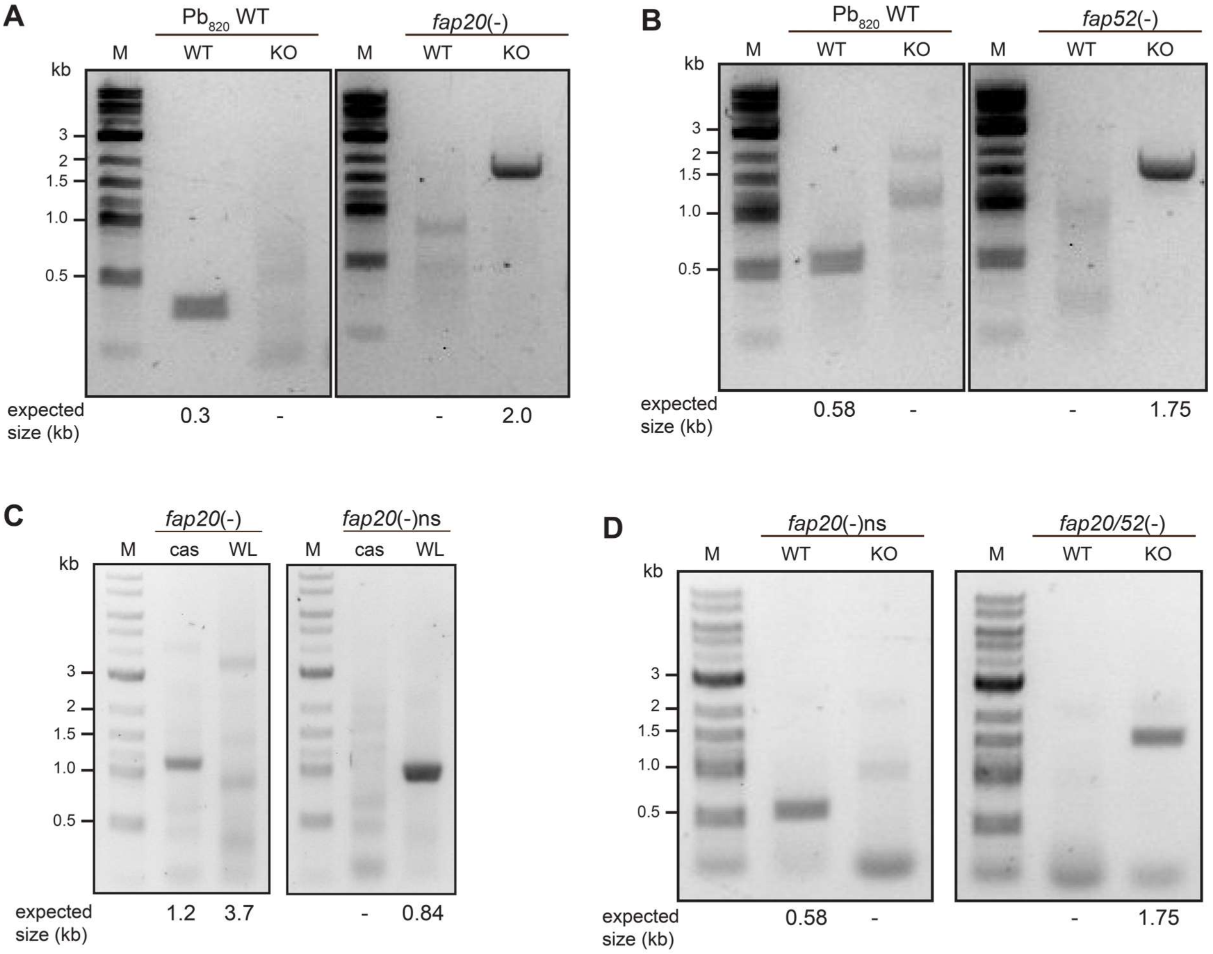


**Supplementary figure 4: Generation of *fap20* and *fap52* single and double knockout parasite lines.** Genotyping PCRs of **A)** *fap20*(-), **B)** *fap52*(-), **C)** *fap20*(-)ns and **D)** *fap20/52*(-). Primers are indicated in supplementary figure 1 and supplementary table 1. Kb, kilobase, ns, negatively selected

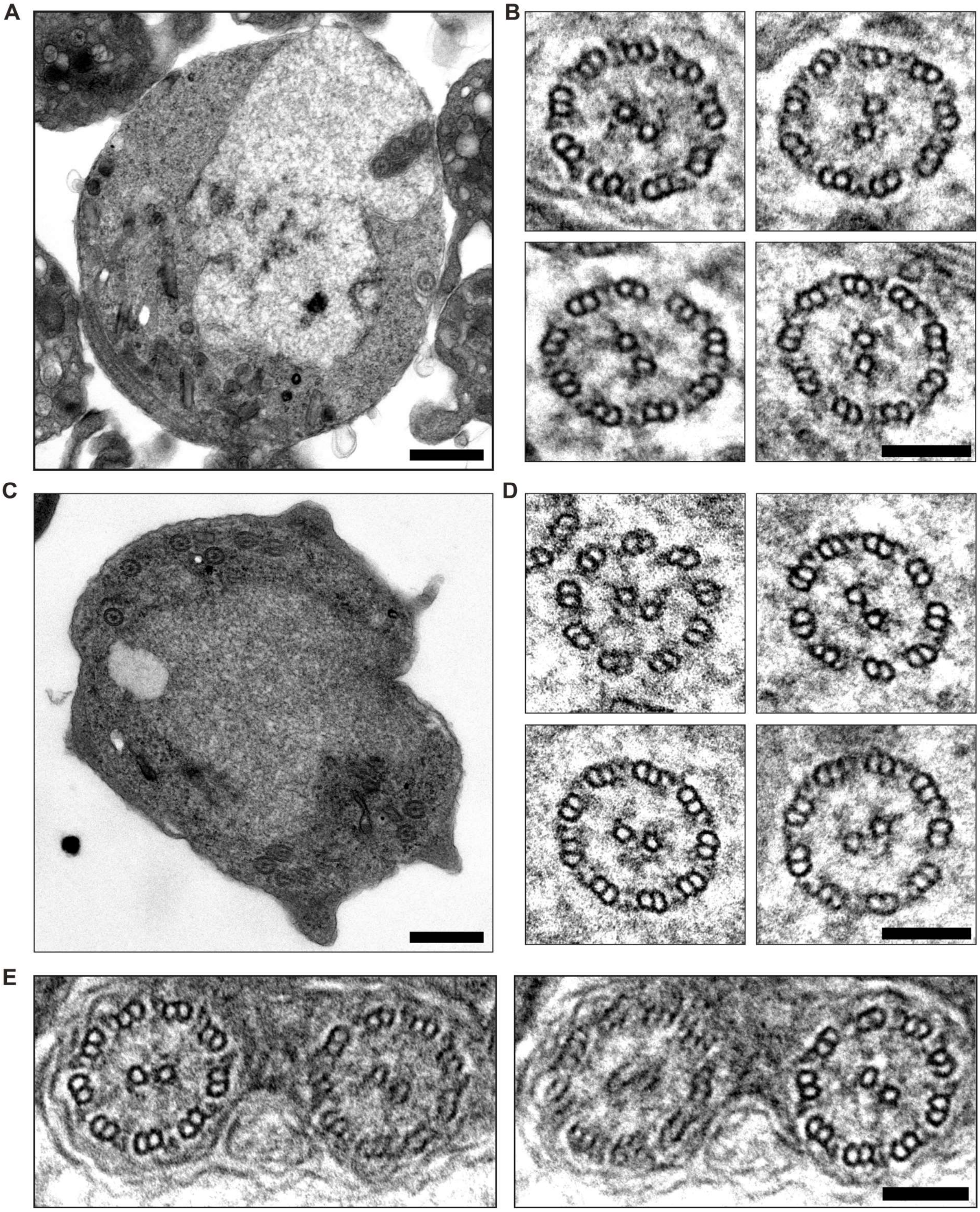


**Supplementary figure 5: Representative TEM images of *fap20*(-) and *fap52*(-) activated gametocytes and axonemes. A, C)** Activated A) *fap20*(-) and B) *fap52*(-) microgametocytes. Scale bar, 500 nm. **B, D)** Close up on B) *fap20*(-) and D) *fap52*(-) axonemes. Scale bar, 100 nm. **E)** Improvement of axoneme orientation via tilting for optimal contrast. Scale bar, 100 nm. Example images from *fap20*(-).

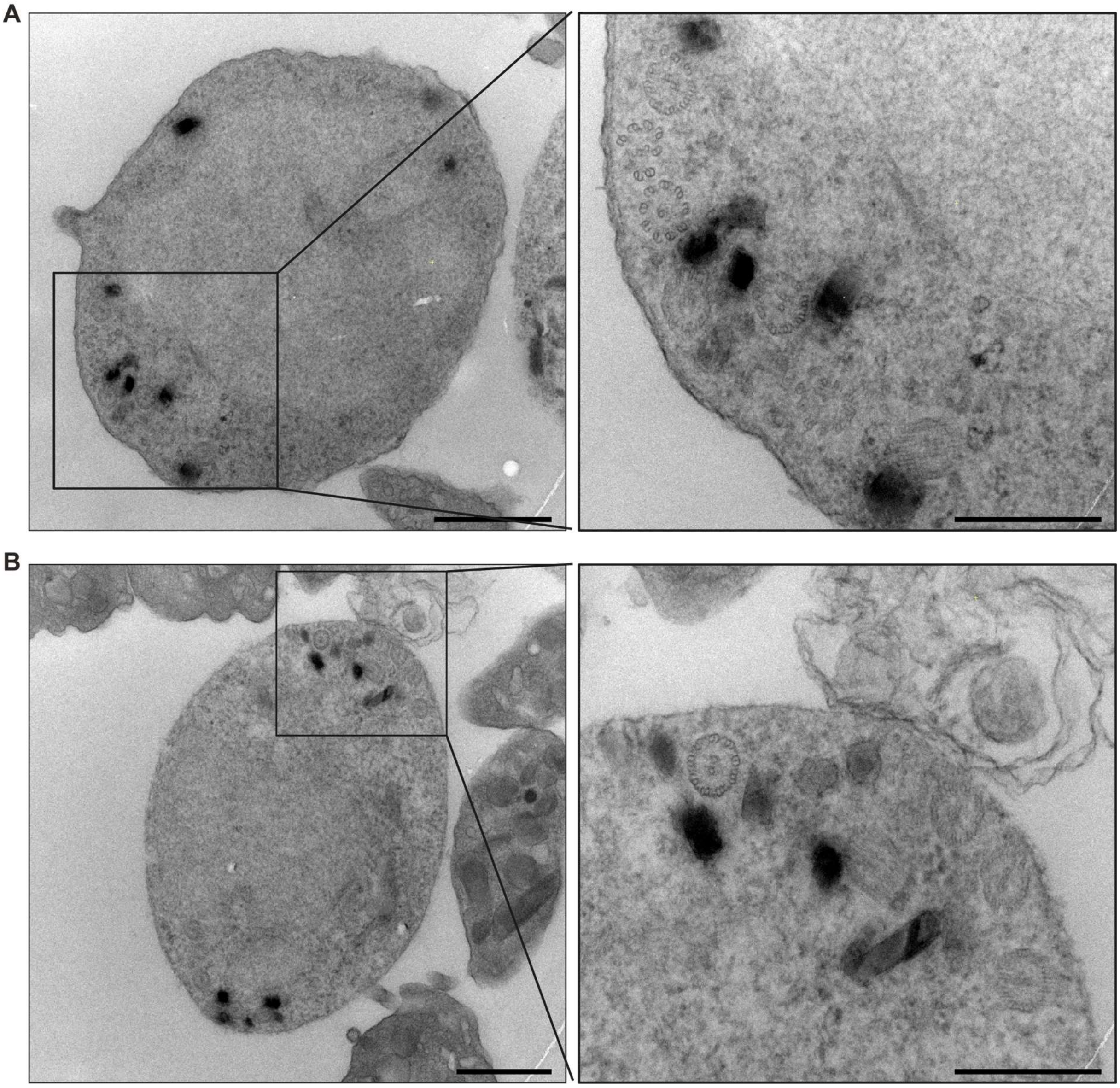


**Supplementary figure 6: TEM images of *fap20/52*(-) male gametocytes**. **A, B)** TEM images of activated *fap20/52*(-) male gametocytes. Region selected for tomogram reconstruction is indicated as box and blow up. Scale bars, 1000 nm (overview) and 500 nm (blow up).

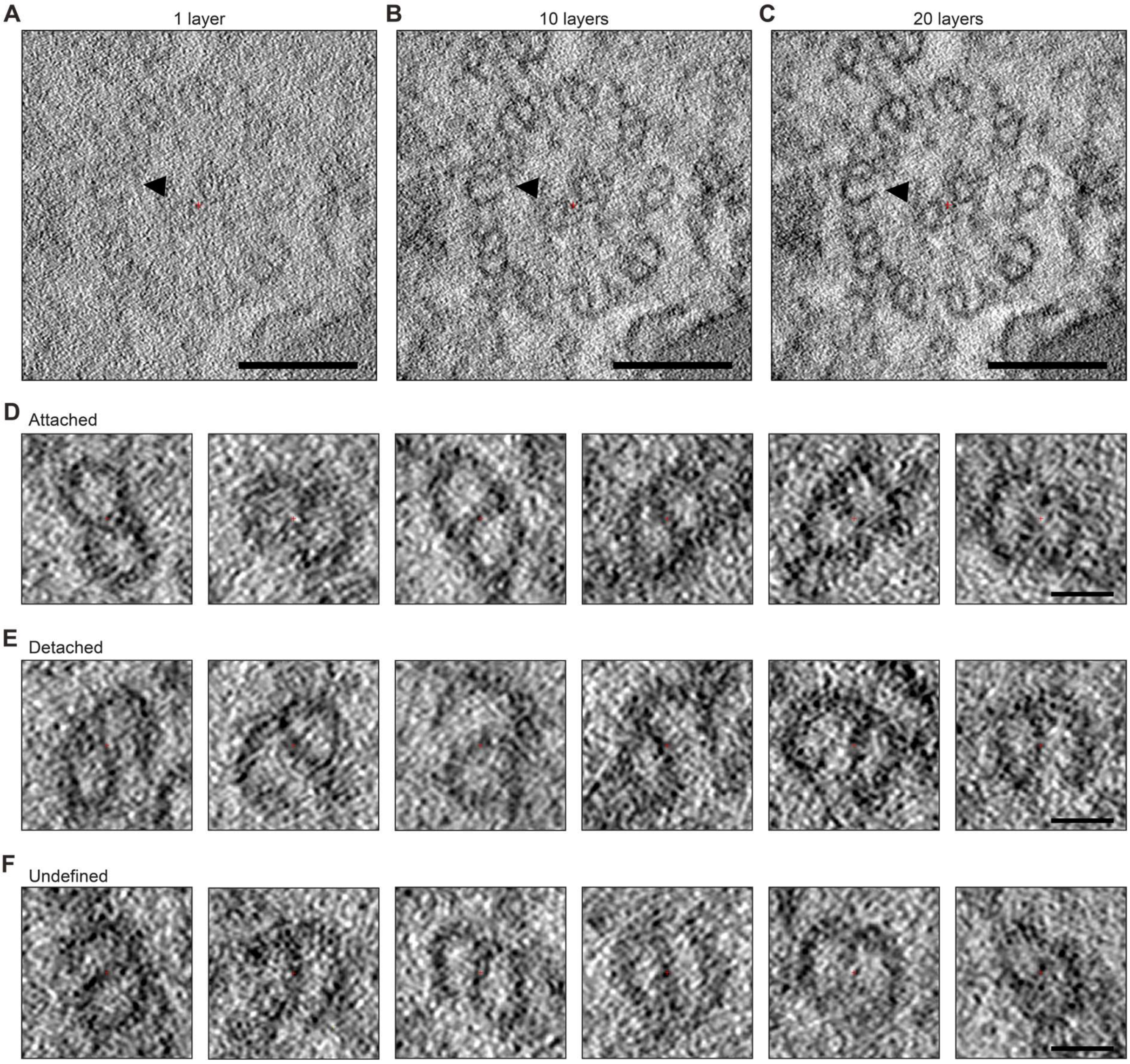


**Supplementary Figure 7: Contrast improvement and exemplary doublet images of *fap20/52*(-) axonemes. A-C)** Greyscale TEM images with sum projection of A) single z layer, B) 10 z layers or C) 20 z layers. Sum projections of 20 z layers were used for all image analysis. Scale bars, 100 nm. **D-E)** Example images of D) attached, E) detached and F) undefined doublets. Scale bars, 20 nm.

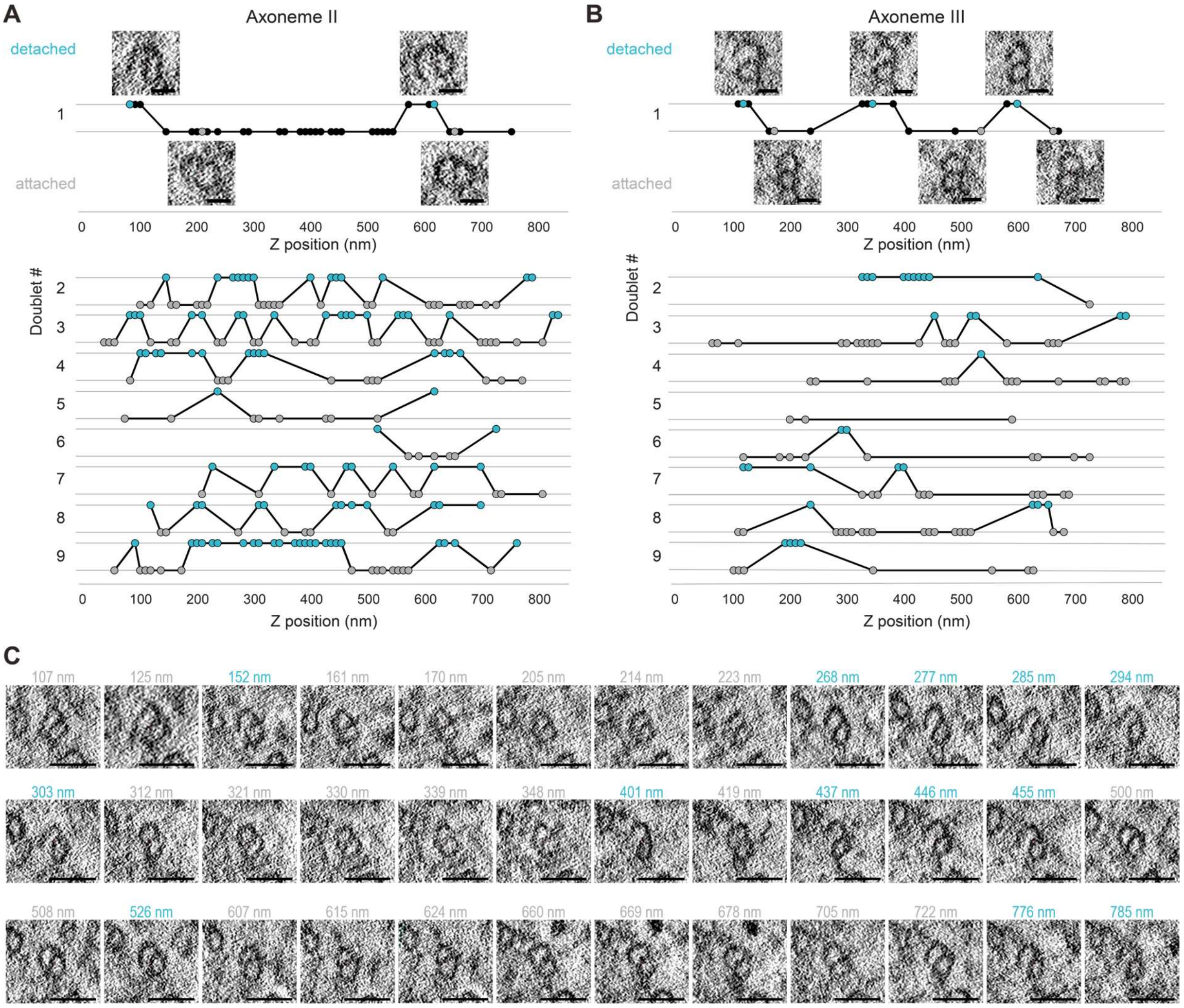


**Supplementary Figure 8: Doublet state across microtubules in *fap20/52*(-). A, B)** Doublet states of doublets 1 to 9 of A) axoneme II and B) axoneme III across z position. Each dot represents an assignable doubled image. Turquoise, detached, grey, attached. First row, example images corresponding to coloured dots are shown, black, data points without example picture. Scale bar, 20 nm. Scale bar, 20 nm. **C**) Images of z slices of axoneme II, doublet 2.

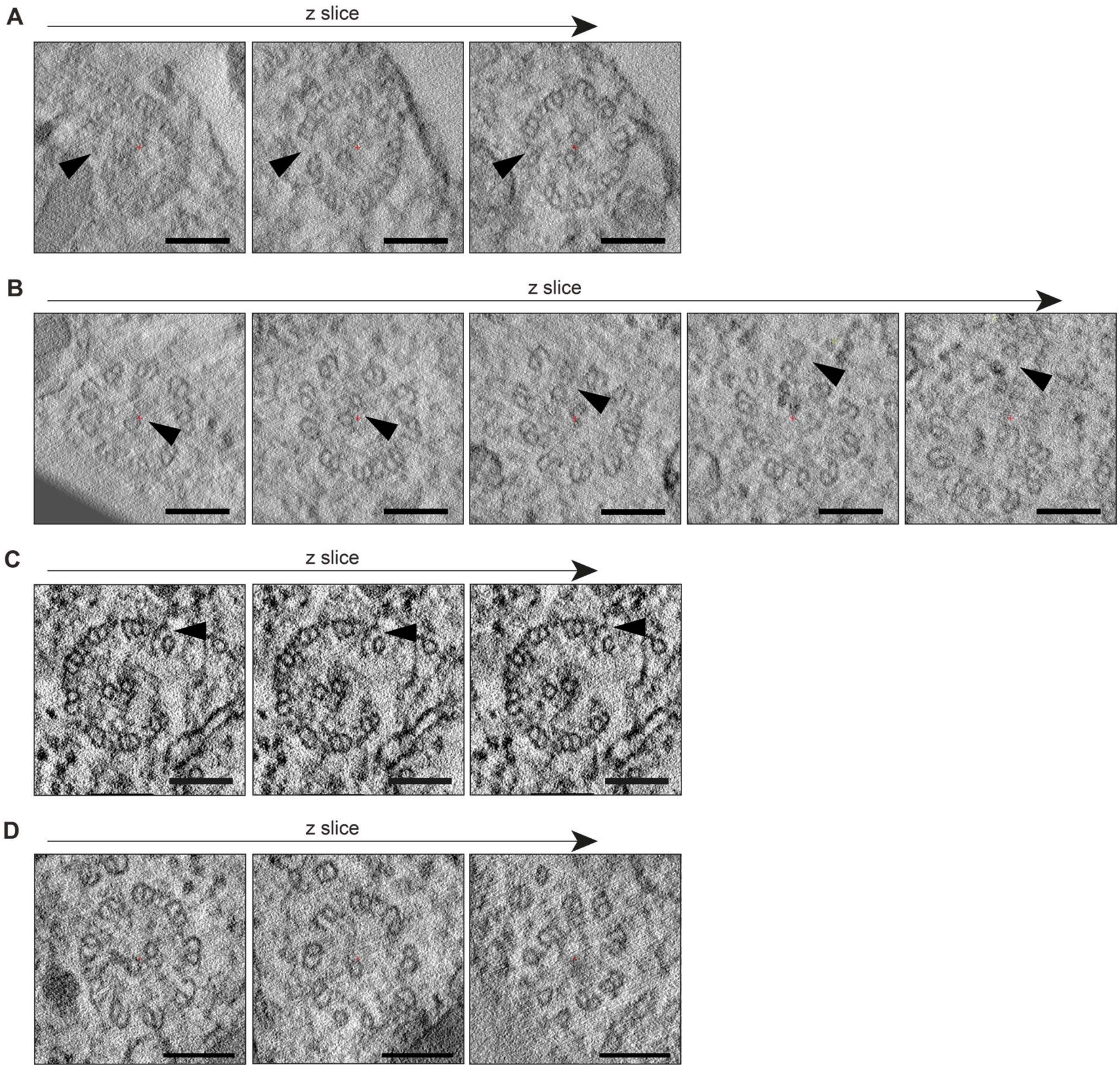


**Supplementary Figure 9: Examples of disordered axoneme arrangement in *fap20/52*(-). A)** The axoneme seems to open at one side (black arrowhead), with doublet MTs moving towards the central pair. **B)** The central pair (black arrowhead) leaves the centre of the axoneme and bends into the cytoplasm of the parasite. **C)** A single doublet (black arrowhead) where both ends of the B tubule detach from the A tubule. **D)** Axonemal tip where central pair disappears and doublets loose arrangement across Z. **A-D)** Scale bars, 100 nm.

**Supplementary Table 1: Primers used for genotyping**

| Genetic modification | PCR | Primer position* | Primer sequence |
|---|---|---|---|
| *spm1*(-)<br>*trxl1*(-)/*spm1*(-) | KO | P4.2 | GAAGAATGATGCTAGCGGAGG |
| | | P2 | ACGTGCATTTCTTAGCGTTTCCT |
| | WT | P3.2 | GTGTGGCTTTCATAGATGCCC |
| | | P2 | TGCCTCTTCCTATGAGACTGA |
| | WL | P1 | CACAACACATAAAAAATGCGCACC |
| | | P2 | ACGTGCATTTCTTAGCGTTTCCT |
| *trxl1*(-) | KO | P2 | ACGTTCTCCACATTGGCAAA |
| | | P4.2 | GATGTCCAGGAGGAGAAAGGC |
| | WT | P2 | ACGTTCTCCACATTGGCAAA |
| | | P3.2 | GGCCCAAGAAGCGATGTCCCA |
| | WL | P2 | ACGTTCTCCACATTGGCAAA |
| | | P1 | AGCGCGCATTAGCCAATTCT |
| *trxl1*(-)/*spm1*(-)ns | KO | P2 | TGCCTCTTCCTATGAGACTGA |
| | | P4.2 | GAAGAATGATGCTAGCGGAGG |
| | WL | P2 | TGCCTCTTCCTATGAGACTGA |
| | | P1 | AAGGCGCATAACGATACCAC |
| *fap20*(-) | WT | P1 | ACCCCGCGCTTAACAAGAGTGA |
| | | P3.1 | TGTACGTACATGTAGGGAAA |
| | KO | P1 | ACCATGGACAGCTAGTTGTGCTCA |
| | | P4.1 | TAATTCAAAGGGACGAGG |
| *fap52*(-)<br>*fap20*(-)/*fap52*(-) | WT | P1 | AGTTGCAGTGAAGACGGGCT |
| | | P3.1 | AGCATCATTGTGGGCATTTCCT |
| | KO | P2 | ACTGATAAAGATTCGCGGTATGGA |
| | | P4.2 | GTAAACTTAAGCATAAAGAGCTCG |
| *fap20*(-)ns | cas | P2 | TGTACGTACATGTAGGGAAA |
| | | P4.2 | TAATTCAAAGGGACGAGG |
| | WL | P2 | TGTACGTACATGTAGGGAAA |
| | | P1 | AAGGCGCATAACGATACCAC |

* Please refer to Supplementary Figure 1 for generic primer binding sites.

2. M. Varadi, S. Anyango, M. Deshpande, S. Nair, C. Natassia, G. Yordanova, D. Yuan, O. Stroe, G. Wood, A. Laydon, A. Žídek, T. Green, K. Tunyasuvunakool, S. Petersen, J. Jumper, E. Clancy, R. Green, A. Vora, M. Lutfi, M. Figurnov, A. Cowie, N. Hobbs, P. Kohli, G. Kleywegt, E. Birney, D. Hassabis, S. Velankar, AlphaFold Protein Structure Database: massively expanding the structural coverage of protein-sequence space with

high-accuracy models. *Nucleic Acids Res.* **50**, D439–D444 (2022).
